# Towards reconstruction of the human interactome from positive and negative experimental evidence

**DOI:** 10.64898/2026.09.09.750397

**Authors:** Joel Ås, Konstantin Pelz, Judith Bernett, Francesco Battini, Markus List, David B. Blumenthal, Martin H. Schaefer

## Abstract

Protein-protein interactions (PPIs) have been detected and reported in the millions, but while they are used in many different contexts for better understanding cellular processes in health and disease, the knowledge of the human PPI network is far from complete, containing many false positive measurements and being highly biased. Both to chart the extent of those problems and to solve them requires not just a knowledge of high-confidence positive interactions, but also likely non-interacting protein pairs. However, this information is typically not reported in PPI studies. We developed a methodology to reconstruct this knowledge from existing PPI data. We reconstruct the experimental search space in which PPI screens have been performed and then create a model that informs how likely a PPI is real given its testing and observation frequency. We argue that negative protein pairs allow us to estimate the error rates of experimental and computational screens. We show how this knowledge could be incorporated for calibration. Finally, we evaluate a simple machine learning approach to PPI prediction and propose how such negative data can be used for training instead of random protein pairs. Together, our results show that reconstructing the experimental search space recovers a largely overlooked layer of information from existing PPI data that can help guide a more accurate and complete mapping of the human interactome.

## Introduction

More than 10,000 [1] protein-protein interaction (PPI) detection studies over the last two decades have uncovered more than one million human PPIs. This knowledge is widely leveraged to understand cellular pathways [2], disease mechanisms [3,4] and the consequences of molecular perturbations [5], and its systematic analysis has given rise to the field of network biology. Large-scale experimental screens using affinity-purification mass spectrometry (AP-MS) [6,7] and yeast two-hybrid (Y2H) [8] assays have produced a substantial fraction of the reported human interactome, but these experiments are costly, labor-intensive, and prone to both false-positive and false-negative detections [6,9,10]. Hence, our knowledge of human PPIs remains incomplete [8] and error-prone. Acknowledging the limitations of PPI detection methods, confidence scores are often applied to quantify the reproducibility of PPIs or the reliability of experimental methods [11,12], thereby selecting interactions considered most likely to be true. While this strategy reduces noise inherent to PPI networks, it can further bias the resulting networks towards well-studied proteins and interactions. Indeed, when PPI networks aggregated across studies are considered, the number of reported interaction partners of a protein is strongly determined by how frequently that protein has been experimentally tested [13]. Due to the high false-positive rates, repeated testing alone increases the probability that an interaction will eventually be reported.

Computational prediction of PPIs offers an attractive complement: once trained, a model can score candidate protein pairs at negligible experimental cost and, in principle, at accuracies approaching or surpassing those of experimental detection methods [14,15]. However, that promise is undercut by how prediction methods are commonly trained and evaluated. Reported performance can be inflated by information leakage between training and evaluation data, including overlapping proteins across data splits and differences in degree distributions between positive and negative examples or other biases [16,17]. When these factors are controlled, measured performance is generally substantially lower. In addition, computational approaches require not only high-confidence positive interactions but also appropriate negative examples. These are commonly generated by sampling from the complement of the known interaction network [18], thereby treating non-reported protein pairs as negatives.

A problem is that interaction databases almost exclusively report positive observations while not explicitly describing the experimental search space in which these observations were made. Consequently, a protein pair detected once in a single experiment is represented in databases in the same way as one detected once following dozens of opportunities for detection, resulting in a compounding of testing and publishing bias. Likewise, a protein pair repeatedly tested but never detected is represented in the same way as a protein pair that has never been tested. Other approaches have aimed at closing this gap before, most notably the Negatome [19]. In their approach they aggregated, manually curated negative information, semantic text-mining and crystal structure data, identifying co-crystalised proteins that were not in direct contact. Already in 2017, Alvarez-Ponce argued for the importance of reporting negative interactions and that this information would not have to be stored explicitly but implicitly be storing which proteins have been tested [20]. Similarly, we argue here that we can partly reconstruct the experimental search space given the reported information: In multi-bait interaction screens, a prey detected with at least one bait must have been present and detectable within that experimental system and therefore defines part of the search space of the other baits tested in the same screen. This principle resembles the logic underlying interaction-calling approaches such as SAINT [21] and ComPASS [22], which evaluate prey signals across multiple baits within individual AP-MS experiments within the same study. However, whereas these approaches operate on the primary quantitative measurements of an individual screen, we use reported binarized interactions to retrospectively reconstruct the search spaces of large numbers of published experiments after their aggregation into PPI databases. A similar idea was presented as viability analysis [23] where they estimated the non-interactions from a single Y2H study including both bait and prey in the searchspace. We apply this principle only from the prey proteins and across 5.530 select AP-MS and Y2H studies, where we identify 97 million distinct tested protein pairs, across 182 million individual tests from approximately half a million reported interactions. By integrating the number of positive observations with the number of inferred tests, we estimate the probability of detection for individual protein pairs and derive sets of high-confidence interactions and non-interactions.

Reconstructing this experimental search space opens several possibilities for improving interactome mapping. Experimentally supported non-interactions provide negative reference sets for training and benchmarking computational PPI predictors, avoiding the assumption that randomly sampled non-reported pairs constitute true negatives. Repeatedly supported positive and negative pairs can provide reference sets for estimating false-positive and false-negative rates and, hence, calibrating experimental detection methods, providing the same function as the human Random Reference Set [10], which provides a reference of non-interaction, to calibrate false positive rates (assumed around 0.1%). We aim to use experimentally validated non-interactions for this purpose instead. And relating positive detections to the underlying testing frequency can reveal assay-specific detection biases and proteins with unusual detection propensities, including sticky proteins [24] in AP-MS approaches or autoactivators in Y2H [8,25,26].

## Results

### Mapping the search space

PPI studies typically do not specifically report the search space. However, we claim that this can be reconstructed by leveraging reported positive interactions. In any multi-bait screen, a prey protein observed at least once likely was present and detectable in that experimental system for all baits, allowing for the reconstruction of the search space of that screen (**Figure 1A**). This inference lets us separate negative data from unobserved or untested protein pairs, while still being easily applicable to aggregated interaction data. Consequently, the number of times a protein pair has been tested versus how many times it has been observed as interacting can be estimated (**Figure 1B**). The estimation provides per-pair detection probabilities and credible intervals from which we define high-replication interactions (HRIs) and high-replication non-interactions (HRNIs). Applying this framework across selected studies curated in IntAct [1], we reconstruct a large experimentally supported non-interaction network.

**Figure 1:**
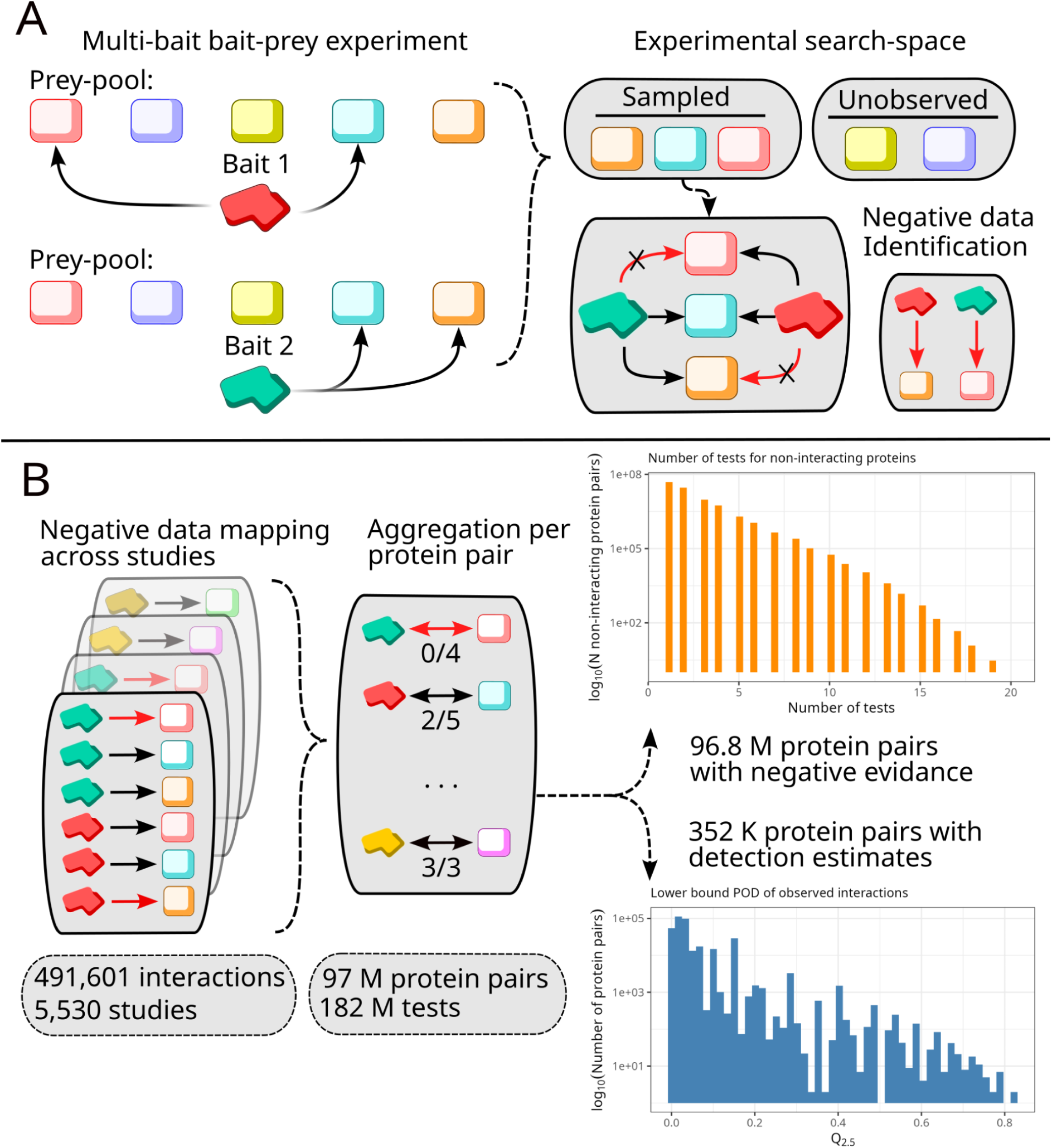
Sampling of experimental search-space and negative data mapping. (A) In a multi-bait experiment, the detectable prey proteins are the same for each bait. Specifically, in AP-MS methods, the prey is dependent on cell-line, protein characteristics and experimental detection setup, but those variables are the same for each bait. Therefore, the detection of a prey from one bait provides a sample of the search space for another bait, providing information on what was tested but not reported as interacting. In turn, this allows for non-interaction data to be inferred from binarized interaction studies. (B) Aggregating the results over multiple studies allows for estimating the number of times a protein pair was reported as interacting versus the number of times it has been tested. Applying this strategy for selected detection methods across the interaction aggregation database IntAct allowed us to map 491,601 interactions across 5,530 studies to a total of 97 M protein pairs tested across 182 M tests. From the replication ratio of interacting proteins, we estimated the lower bound probability of detection of a protein pair (*Q*_2.5_, see Methods) and identified 96.8 M protein pairs with experimental evidence of non-interaction.

From the 491,601interactions reported in the selected studies, we inferred the protein pair replication rates (**Figure 1B**), resulting in 97 million unique tested protein pairs spanning 182 million individual tests. These pairs were not tested uniformly (**Figure 1B**): A small number were probed multiple times across studies, while most were tested only once or twice.

This testing inequality allows us to call interactions and non-interaction with graded confidence, as a tested pair’s credibility increases with reproducibility. To model the credibility of the results, we considered the probability of detection (POD) as our ranking metric. For each protein pair we modelled the number of detected interactions and tests using a beta-binomial model. The beta prior were set to reflect the global detection rate of all tests performed. This model provides us with a posterior distribution of where the true probability of detection might lie. From the posterior distribution we can get the the 95% credible interval contained between *Q*_2.5_ and *Q*_97.5_. These estimates limits represents the section where the POD-value is 97.5% more likely to be above the estimate (*Q*_2.5_) or below the estimate (*Q*_97.5_). Essentially, this allows us to consider a protein pair and evaluate its status as an interaction on *Q*_2.5_ and its status as non-interactor on *Q*_97.5_ (see Methods). To define a set of HRIs we filtered the full dataset on *Q*_2.5_ > 0. 29. This threshold is chosen as a reasonable trade-off between detection ratio and number of tests. At 0.29 a pair needs to be replicated as interacting 3 out of 3 times while a pair at detection ratio ∼0.7 needs at least 5 out of 7 tests succeeding and as the detection ratio decreases the minimum number of tests increases (24 required tests at a ratio of 0.5). This provided us with a set containing 6,221 interactions (mean detection ratio= 0.934, the most tested pair was tested 69).

Due to the very low detection prior (**Table 1**) a protein pair reported as interacting once retains a *Q*_97.5_ above the untested baseline probability until it has been tested more than 1,383 times. The most tested pair in our data has been tested 69 times, therefore any protein pair with any detection cannot exist in our HRNI set. The remaining tested protein pairs has therefore no observed interactions, resulting in a monotonically decreasing *Q*_97.5_ with the number of tests. Therefore we defined the HRNI based on the minimum number of test (without observed interactions) instead of thresholding *Q*_97.5_. Setting the same number of minimum tests of three tests for symmetry of the HRI threshold. The resulting HRNI set contains 18,982,508 protein pairs (average tested = 3.9;max number of tests = 20).

**Table 1:** Observed interactions, total tests, and detection ratio per dataset.

| Dataset | Proteins pair with observed interactions | Total experiments | Alpha value for prior | Beta-value for prior (1- alpha) |
| --- | --- | --- | --- | --- |
| MS | 355,406 | 135,279,697 | 0.002627 | 0.997373 |
| Y2H | 124,195 | 46,582,773 | 0.002666 | 0.997334 |
| Combined (Y2H+MS) | 479,601 | 181,862,470 | 0.002637 | 0.997363 |

We made the negative protein pairs tested five or more times available on https://hippie-db.net.

### Validating non-interacting pairs

We next aimed to functionally validate our newly defined negative interactions. Interactions, contrary to their absence, are expected to follow guilt by association, a cornerstone of biological-network analysis [27]. Guilt by association implies that interacting proteins tend to share properties, such as participating in the same biological process [28] or predominantly sharing the same subcellular compartments [29]. Naturally, this should hold true for when comparing HRIs to HRNIs. To evaluate this hypothesis, we compared the functional annotation and subcellular localization [30]. We restricted the annotations to biological-process (BP) GO terms [31] and localisation terms annotating 400 or more genes (**Table S2**). This threshold was to limit the number of eligible terms and reduce the influence of multiple-hypothesis testing. For each term, we computed the HRI-versus-HRNI odds-ratio (OR) of a shared annotation. However, annotation is a property of a gene and not protein pairs therefore we estimated the 95 % CI for each annotation bootstrap by resampling shared annotation ratios of proteins (**see Methods, Table S3**) for each annotation. Furthermore, we can estimate if these properties are equal depending on the experimental detection category (**Table 1**).

The HRI pairs shared annotation substantially more often than HRNI pairs (**Figure 2A)**. For the MS and combined datasets, the OR exceeded one and rejected the null of OR = 1 for 24 of 25 terms across both categories (q < 0.05, Benjamini-Hochberg adjusted; **Table S3**). The Y2H dataset showed the same direction but a weaker signal, rejecting the null with OR > 1 for only 9 of 24 terms, and nucleoplasm was the sole localisation over-represented among Y2H interactions.Thus, non-interacting pairs overall lack the functional and spatial co-membership characteristic of interactors, most clearly demonstrated for the AP-MS and combined datasets. The two expectations are also informative: This localisation preference of Y2H is inherent to the method as both proteins must be present in the nucleus in order to activate the reporting phenotype [32]. The single shared not enriched GO-term is proteolysis. Proteolysis consists of protease pathways, whose role is to bind and digest proteins. While this would lead to a large number of protein interactions, the annotation in that interaction is one-sided as it is an interaction between an enzyme and its substrate.

**Figure 2:**
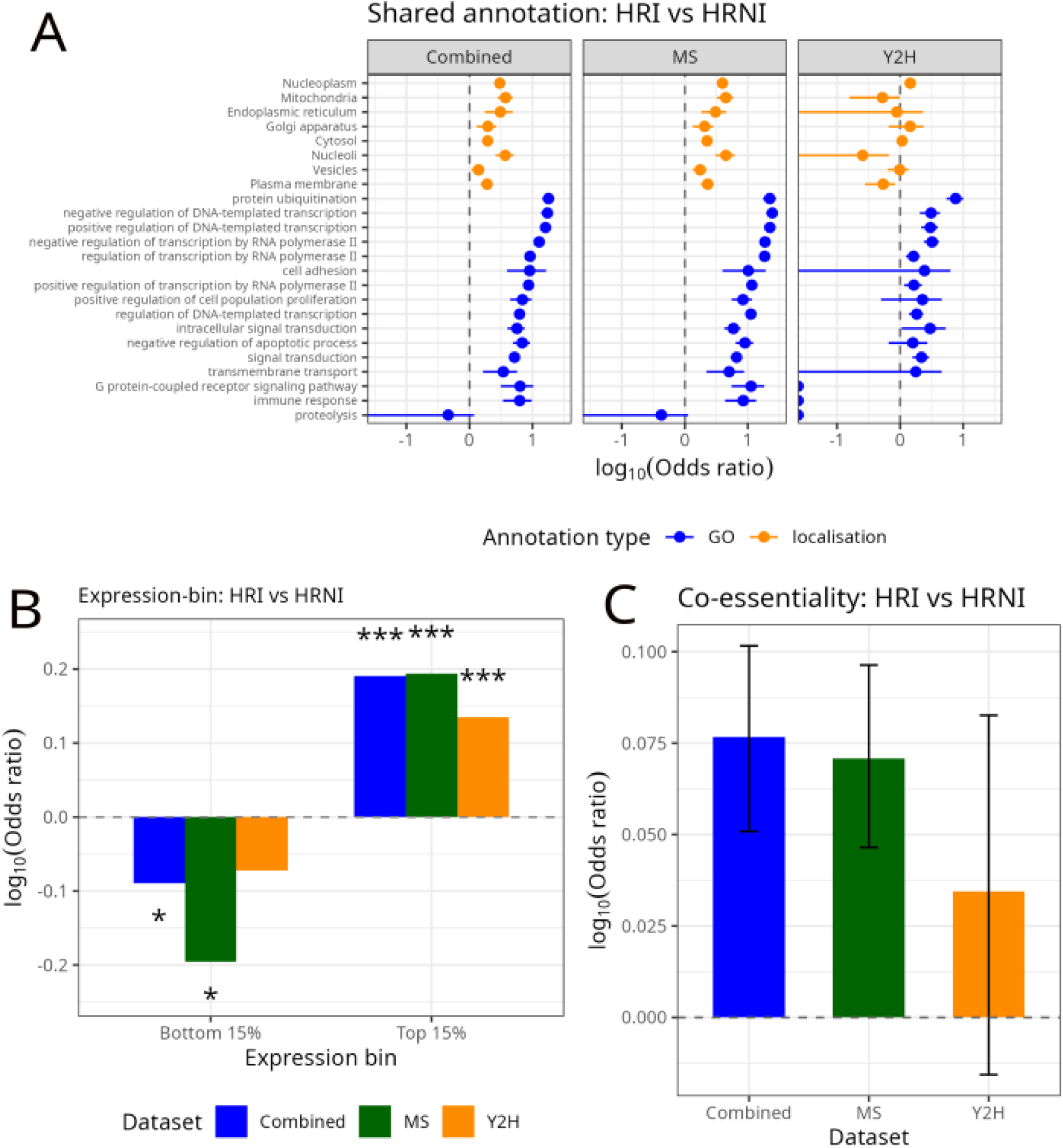
Function, spatial, expression and essentiality is shared among HRI but not among HRNI: In all panels the comparison between degree-balanced high-replication interaction sets (HRI, *Q*_2.5_ > 0.29) versus high-replication non-interaction sets (HRNI, *n_tested_* ≥ 3, *n_obs_* = 0) are reported in 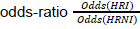. Any error bars are 95% CI intervals bootstrapped across 5000 samplings on a protein basis and as the properties are properties of the constituting proteins and not protein pairs, except co-expression. Each property was compared across three datasets Y2H, MS and Combined (**Table 2**). (A) OR of shared-annotation of subscellular main localisation from Human Protein Atlas and GO biological process terms. The terms tested constituted 400 or more genes to ensure sufficient coverage and reduce the number of hypotheses. The majority of terms were overrepresented for MS and Combined while only Nucleoplasm was overrepresented for Y2H, in line with the nature of the detection method. (B) ARCHS4 pairwise Pearson correlations for co-expression were binned into top 15% and bottom 15%. High correlation was overrepresented among HRIs and low correlation for most of HRNIs (Fisher’s exact test; asterisks mark p < 0.05, p < 0.01 and p < 0.001). (C) Gene 90 % confidence loss of function tolerance (LOEUF) were obtained from gnomad v2.1.1. The 25% bottom percentile across all genes were estimated and labeled as intolerant and shared intolerant status tested, showing moderate but significant enrichment for HRIs among MS and Combined.

**Table 2:**
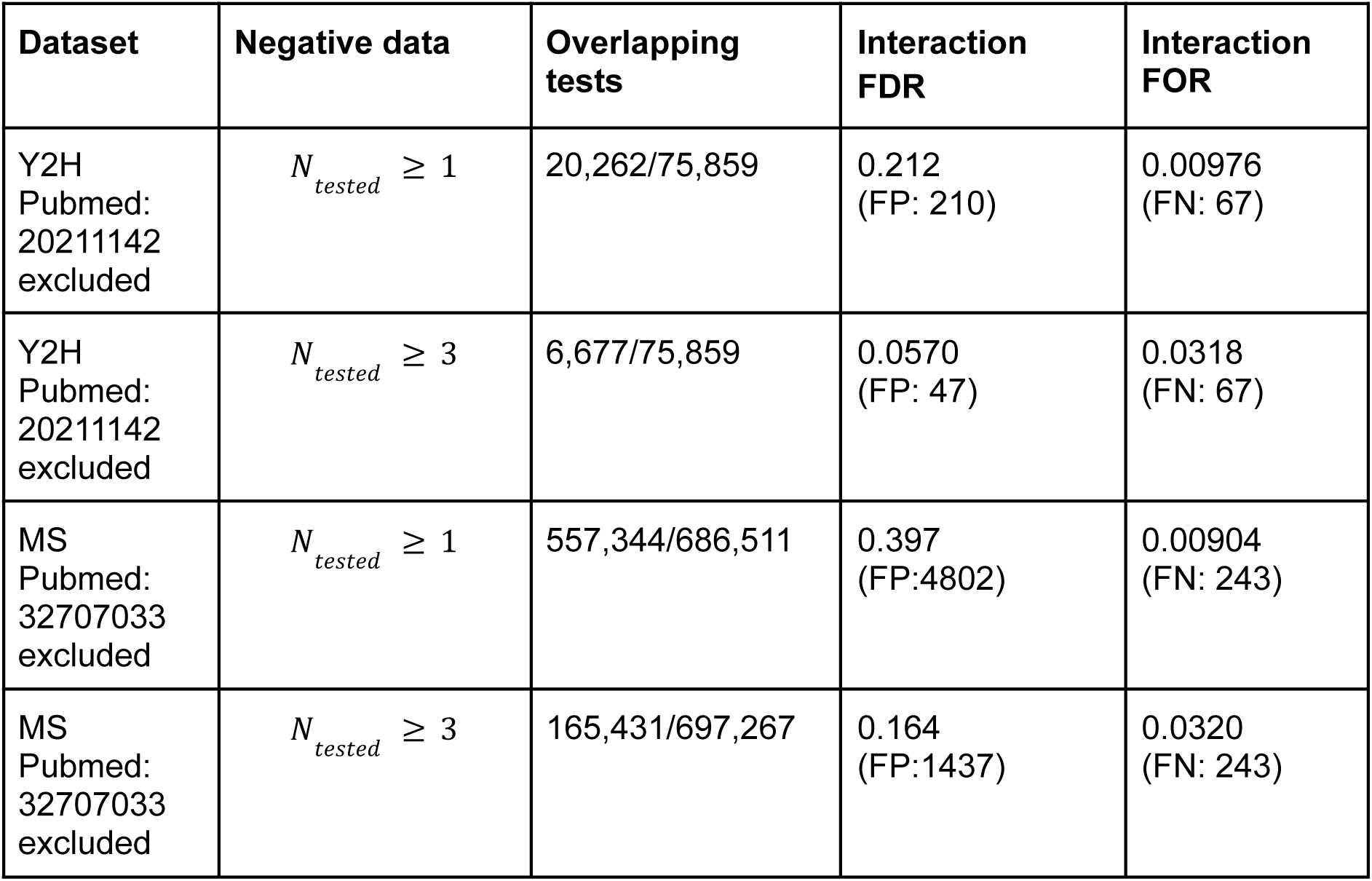
Leave-one-out estimation of false discovery rate (FDR) and false omission rate (FOR), when considering negative data once seen or thrice replicated. Positive reference data is any previous seen interaction.

| Dataset | Negative data | Overlapping tests | Interaction FDR | Interaction FOR |
| --- | --- | --- | --- | --- |
| Y2H<br>Pubmed:<br>20211142<br>excluded | $N_{tested} \geq 1$ | 20,262/75,859 | 0.212<br>(FP: 210) | 0.00976<br>(FN: 67) |
| Y2H<br>Pubmed:<br>20211142<br>excluded | $N_{tested} \geq 3$ | 6,677/75,859 | 0.0570<br>(FP: 47) | 0.0318<br>(FN: 67) |
| MS<br>Pubmed:<br>32707033<br>excluded | $N_{tested} \geq 1$ | 557,344/686,511 | 0.397<br>(FP:4802) | 0.00904<br>(FN: 243) |
| MS<br>Pubmed:<br>32707033<br>excluded | $N_{tested} \geq 3$ | 165,431/697,267 | 0.164<br>(FP:1437) | 0.0320<br>(FN: 243) |

To further explore what we expect to be true among interacting pairs compared to non-interactions, we evaluated co-expression and co-essentiality. For two proteins to interact, both proteins need to be present and expressed. Though not necessarily highly co-expressed, there should be an enrichment of high co-expression among interactors compared to non-interactions. To test this, we used the precomputed pairwise Pearson correlations of expression data from ARCHS4 [33], a resource providing processed RNA-seq obtained from the functional genomics database Gene Expression Omnibus [34]. To annotate the data for later testing, we considered the full co-expression density and labelled any protein pair in the bottom 15% as “low” or top 15% “high. Co-expression is an attribute of the edges (not protein as with annotation), therefore instead opted to degree-balance HRNI and HRI sets (protein specific degree spearman rho:. 97 MS; 0.98 Y2H, 0.97 Combined. **Method S1**), this means each protein has equal representation of edges in both sets. Due to the balance and the edge-attributes we could instead use a Fisher’s exact test to establish over– or under-representation among binned co-expressions (**Figure 2B**). In accordance with the hypothesis, high coexpression was enriched among HRIs for all datasets [OR: 1.56 – 1.36]. Low co-expression was significantly enriched among HRNI for the AP-MS [OR: 0.64] and the combined dataset. This result can be explained by fundamental differences between AP-MS and Y2H testing: In AP-MS, the bait is transduced while prey are endogenous, so a prey that is poorly co-expressed with the bait is less likely to be detectable in that experimental system. In Y2H, both partners are expressed artificially in yeast, making detection independent of human cell co-expression.

Furthermore, guilt-by-association extends to protein essentiality. If two proteins form complexes or partake in pathways, loss of function impacts that biological function. Therefore, an interaction involving an essential protein should extend that essentiality to its interaction partners. For estimation of loss-of-function (LoF) tolerance of genes, we used gnomeAD 2.1.1 LOEUF [35], which estimates a gene’s LoF tolerance across human genomes with a 90% confidence interval. We labeled a gene’s LoF tolerance as “*low*” if its LOEUF estimate were below the 25% quantile of all estimates (of all human genes). Shared LoF tolerance bin was tested on degree-balanced HRI and HRNI sets in a similar manner as co-annotation: OR estimates with protein-annotation rates bootstrapped 5000 times with resampling. Pairs in which both proteins are LoF-intolerant were modestly but consistently enriched among HRIs relative to HRNIs (**Figure 2C**) except for Y2H. The enrichment were clearest in the AP-MS-derived data (OR 1.18, 95 % CI 1.11–1.25, n=4171, p=2e-4) and in the combined set (OR 1.19, 95 % CI 1.12–1.26, n=5350, p=2e-4), while the Y2H estimate did not reach significance (OR 1.08, 95 % CI 0.96–1.21, n=2617, p=0.16).

### Experimental negatives as a benchmarking tool

After establishing that non-interactions behave in the expected way, we ask if we can use them for the applications mentioned above. The first application evaluated is its potential as a benchmarking tool against future and current experimental screens. In both AP-MS and Y2H detection screens, precision is paramount as faulty error estimations can lead to inclusions of erroneous interactions. While positive controls are available, such as the human positive reference sets [36,37], the lack of experimentally supported non-interactions makes it difficult to estimate accuracy of the panel.

As a proof-of-concept, we excluded a large study from the AP-MS dataset and the Y2H dataset, respectively. These studies were only chosen due to their large relative size to the rest of the data. Then, we retrieved HRIs and HRNIs based on the remaining combined AP-MS and Y2H dataset, and used these to estimate FPRs and FORs for the two excluded studies. We defined two sets with negative data for estimation of false discovery rate and false omission rate – one lenient(*N_tested_* ≥ 1) and one strict (*N_tested_* ≥ 3). For positive data, we consider any observed interaction as true (**Table 2**). Treating any test as a proof of non-interaction we estimate the FDR to be 21% (Y2H) and 40%(MS). With higher stringency on the negative data we observe lower but still significant FDR values of 5%(Y2H) and 16%(MS).

A similar problem exists for computational PPI prediction. The two main branches of these predictors are sequence-based models [38] and structure-informed models [39]. For structure-informed models, AlphaFold-multimer [40] and RF2-PPI [15] are some of the most sophisticated interaction predictors available. However, a protein pair for which no experimentally determined complex structure exists is simply a pair nobody has yet solved, so an absence from structural records reflects the lack of experiments as much as the pair’s capacity to bind. Negative data from bait-prey studies is also an absence of evidence of interaction, but the tests can be aggregated and evaluated based on how often it has been tested. Therefore, we argue that this data would be a good fit to compare to the predicted structure-based interaction results.

Unfortunately, a computational proteome-wide structure-informed interaction screen is outside of our computational budget. We therefore used the published RF2-PPI results for Zhang *et al.’s* data-publication [41]. They applied RF2-PPI across 43.6 M protein-pairs. We estimated the false-discovery rate (FDR) as the fraction of HRNIs over the number of pairs with any reported interactions(n=351,552) (**Figure 3**). We applied three stringency tiers to our HRNI: N_tested ≥ 3 (n = 6,321,239), ≥ 5 (n = 1,530,969) and ≥ 7 (n = 386,143) and estimated FDR above two RF2-PPI prediction probability cutoffs: p>0.5 and p>0.99. For p>0.5, we obtained FDR = 0.886, FDR = 0.651 and FDR = 0.320 across tiers; for p>0.99, we obtained FDR = 0.153, FDR = 0.038 and FDR = 0.010. We thus consistently estimate lower FDRs for high-confidence than for low-confidence RF2-PPI predictions, which provides experimentally supported evidence for the credibility of RF2-PPI’s structure-based predictions.

**Figure 3:**
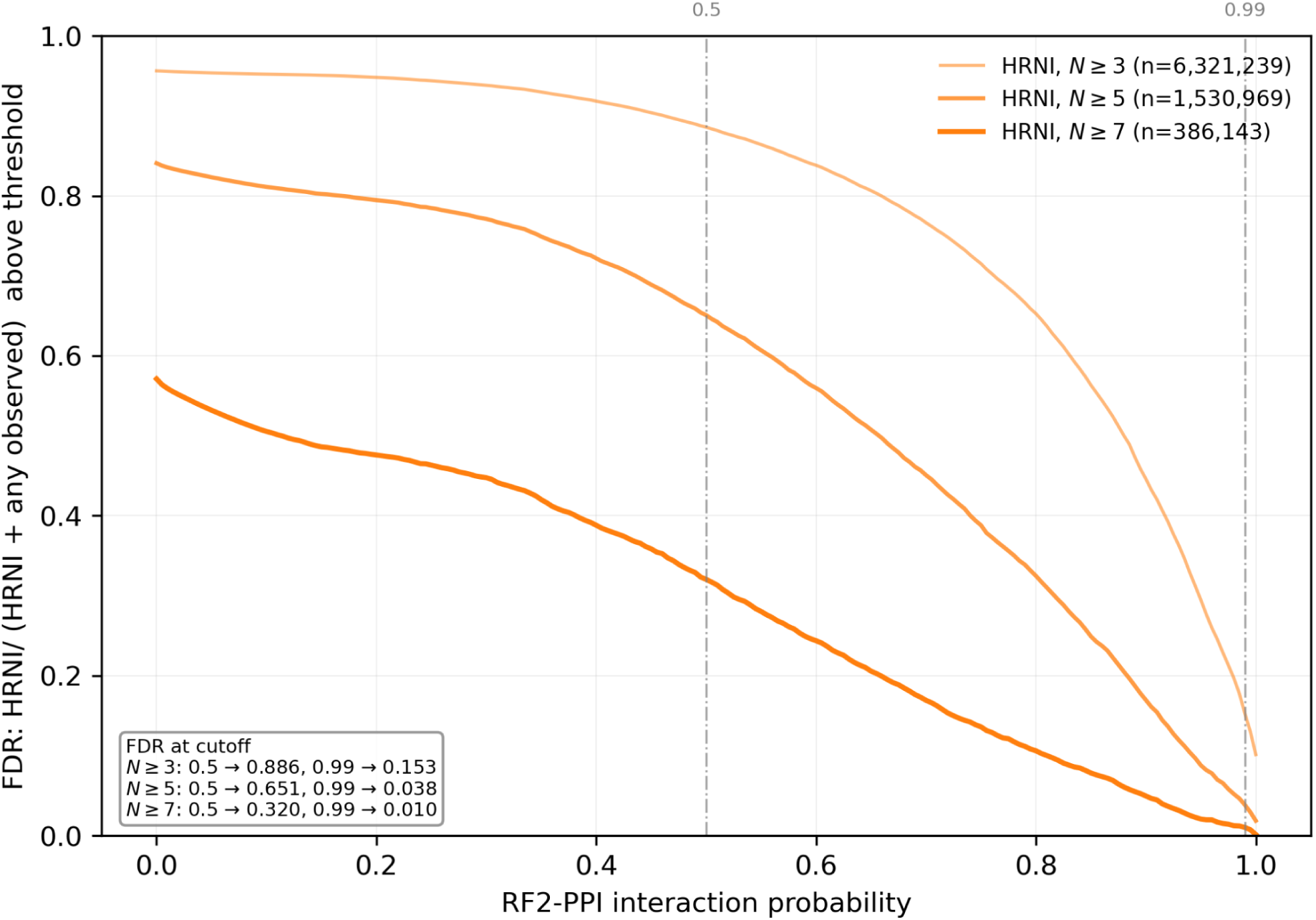
FDR estimates vs RF2-ppi. RF2-PPI probabilities for the 43.6 M pairs preselected on DCA > 0.12 were intersected with a reference set of reported interactions (n = 351,552) and HRNIs at three stringency tiers (N_tested ≥ 3, n = 6,321,239; ≥ 5, n = 1,530,969; ≥ 7, n = 386,143). At each probability cutoff, FDR is the proportion of reference pairs above the cutoff that are HRNI. FDR falls with cutoff for every tier, but the differences between tiers track the size of the HRNI set relative to the interaction set: the corresponding likelihood ratios, FDR/(1 − FDR) × N_obs/N_HRNI, are near-identical across tiers (0.432/0.428/0.428 at p > 0.5; 0.0100/0.0091/0.0092 at p > 0.99).

The FDRs values are influenced by the relative size difference between any observed interaction and HRNI set where the most permissive is 18 times larger than the interaction set. To factor out this relationship we calculated the enrichment as *e* = *N_HRNI_* /*N_obs_* ( *FDR*/(1 − *FDR*))^−1^. Which tells how much more common detections are above p. The enrichment per HRNI were respectively e=2.31, e=2.33 and e=2.33 at p > 0.5 and e=99.54, e=110.25 and e=108.74 at p > 0.99. Meaning that HRNI are roughly as believable when reported in three experiments as in seven.

### Role-specific binding variance ranks auto-activators and contaminants

A central problem of Y2H and AP-MS are auto-activators and sticky proteins. In Y2H this would mean a bait fused with the DNA-binding domain facilitates the activation of the reporter gene independently. This process of auto-activation has been reported for a number of proteins in different experimental setups. During affinity-purification contaminants, or “sticky” proteins, are the proteins that remain detectable after purification, even if the protein is not necessarily bound to the bait, the CRAPome [24] tracks such sticky proteins as a quality control resource for AP-MS workflows. This could be due to a number of factors, such as prey abundance [42], unspecific binding [43] or ease of detection [44]. However, one commonality of sticky proteins and auto-activators in binarized interaction data is that interactions are reported, regardless of the identity of the partnering bait (MS) or prey (Y2H). Meaning that the detection ratio of that protein would be uniform. With negative results, we can estimate the specificity of binding of proteins in either role as bait or prey. Therefore, we selected all protein pairs tested twice or more and estimated the global detection ratio of per bait or prey. Given this detection ratio we could compute the Pearson Chi distance for each protein pair compared to the expected number of detections given the global detection ratio.

From that we could calculate the mean, 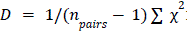: where D is the dispersion between the observed number of detections and the expected detections under a uniform distribution. This implies that a D close or below 1 agrees with a uniform distribution of detection or that the partner protein has little or no effect on detection. The higher D, the more specific its detected binding is (**Figure 4**). To support this model, we compiled a list of known Y2H auto-activators(n=71) and selected a subset of proteins from CRAPome with a mean spectral count > 1 (n=576, among the N replication in CRAPome 1.0). These known problematic proteins tend towards low D-values, many with a high global detectability. The fact that these proteins are known problematic means that they are routinely filtered out in modern interaction workflows. The aggregated data we are working on is a collection of interaction studies spanning decades and multiple studies. As these results show, we are still able to identify multiple flagged problematic proteins through their detection specificity, which can be computed knowing what search-space they have been tested in. Furthermore, this provides a way to identify previously unlabeled auto-activators in Y2H or contaminations in MS-experiment, providing a stickiness value for the proteins.

**Figure 4:**
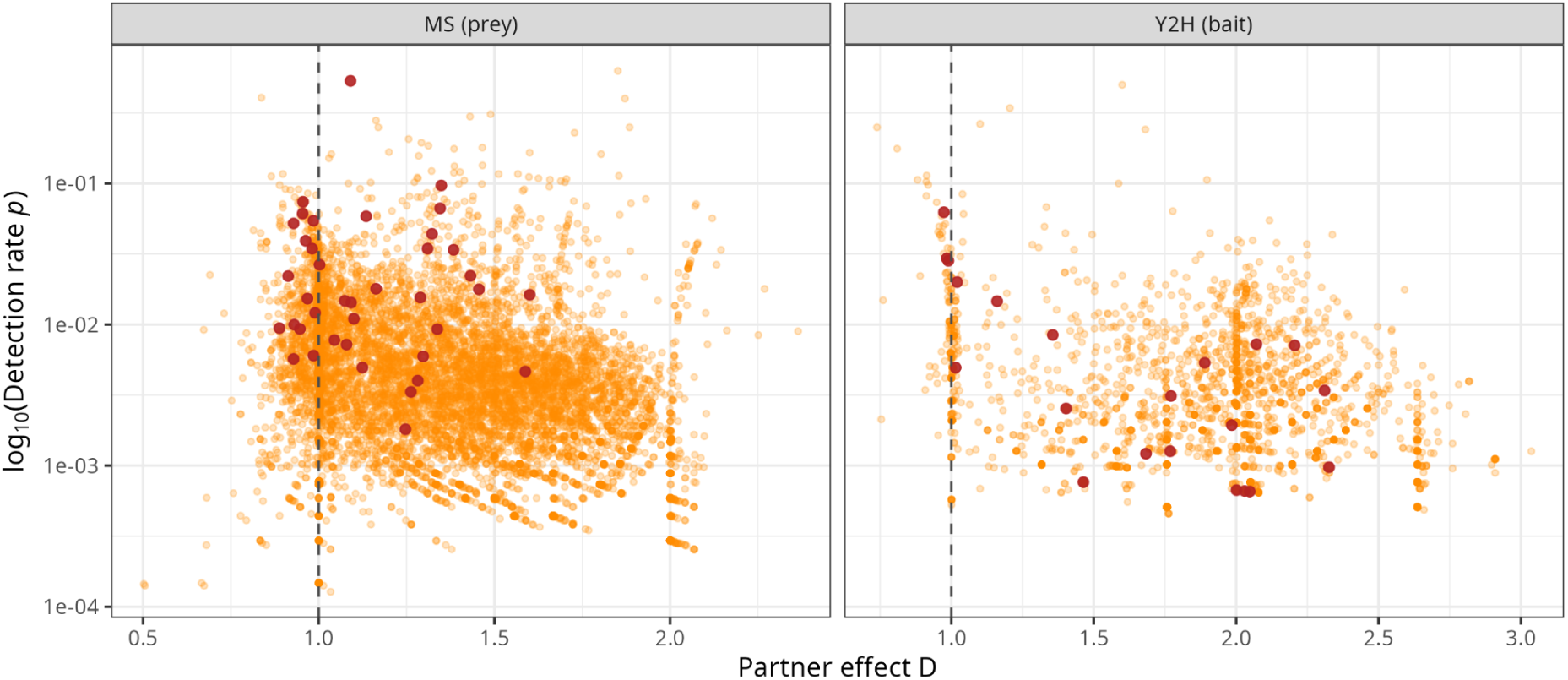
Known auto-activators and sticky proteins detect independently of their partner. For each protein tested in at least two pairs, the expected detections per pair were computed from that protein’s global detection ratio in the given role, and D is the mean Pearson χ² contribution over its pairs. D ≈ 1 indicates detection close to the partner-independent expectation, while larger D indicates partner-specific detection. Each point is one protein, plotted against its global detection rate *p* (log scale); the dashed line marks D = 1. Left, proteins as prey in the MS dataset; right, proteins as bait in the Y2H dataset. Dark red points mark known problematic proteins, CRAPome 1.0 entries with mean spectral count > 1 (n = 576) for MS, and curated Y2H auto-activators (n = 71) for Y2H. In both assays the labelled proteins with the highest detection rates concentrate at D ≈ 1, the region expected for partner-independent detection, whereas the bulk of proteins at comparable detection rates spread to higher D.

### Estimation of the observed interactome and degree distribution

Given a representative search-space for a protein, the detection ratio in that space should agree with that protein’s degree in the true human interactome. Therefore, a sizable enough sample of human proteins would be informative for the interactome’s degree-distribution.

Blumenthal *et al 2024* [13] showed that the observed degree distribution of interactors is bound to be power-law distributed already due to the power-law nature of tested baits. However, we believe that it should be possible to estimate a protein’s degree by knowing how large its search-space has been, not how many times it has been tested as a bait.

First to estimate a protein general detectability, we adjusted for any bait/prey related effects estimating the increase in detectability when a protein is being tested as a bait or prey. In order to have a better estimate of how the proteins act at either bait or prey, select all protein pairs tested at least twice in each orientation. Ensuring that the role effect is estimated within pairs rather across differently composed sets of proteins. Given this subset we estimated the bait and prey detectability of each protein in Y2H and MS (**Figure 5A**). We could observe a moderate correlation of 0.422 for MS and 0.195 for Y2H. Additionally, a general prey detection preference (log2 FC, bait/prey: –0.252) for MS and bait detection preference for Y2H (log2 FC, bait/prey: 0.614) was detected. As we expect the interactions to be undirected in the cell we calculated the adjusted average detection as 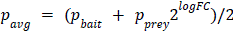

**Figure 5:**
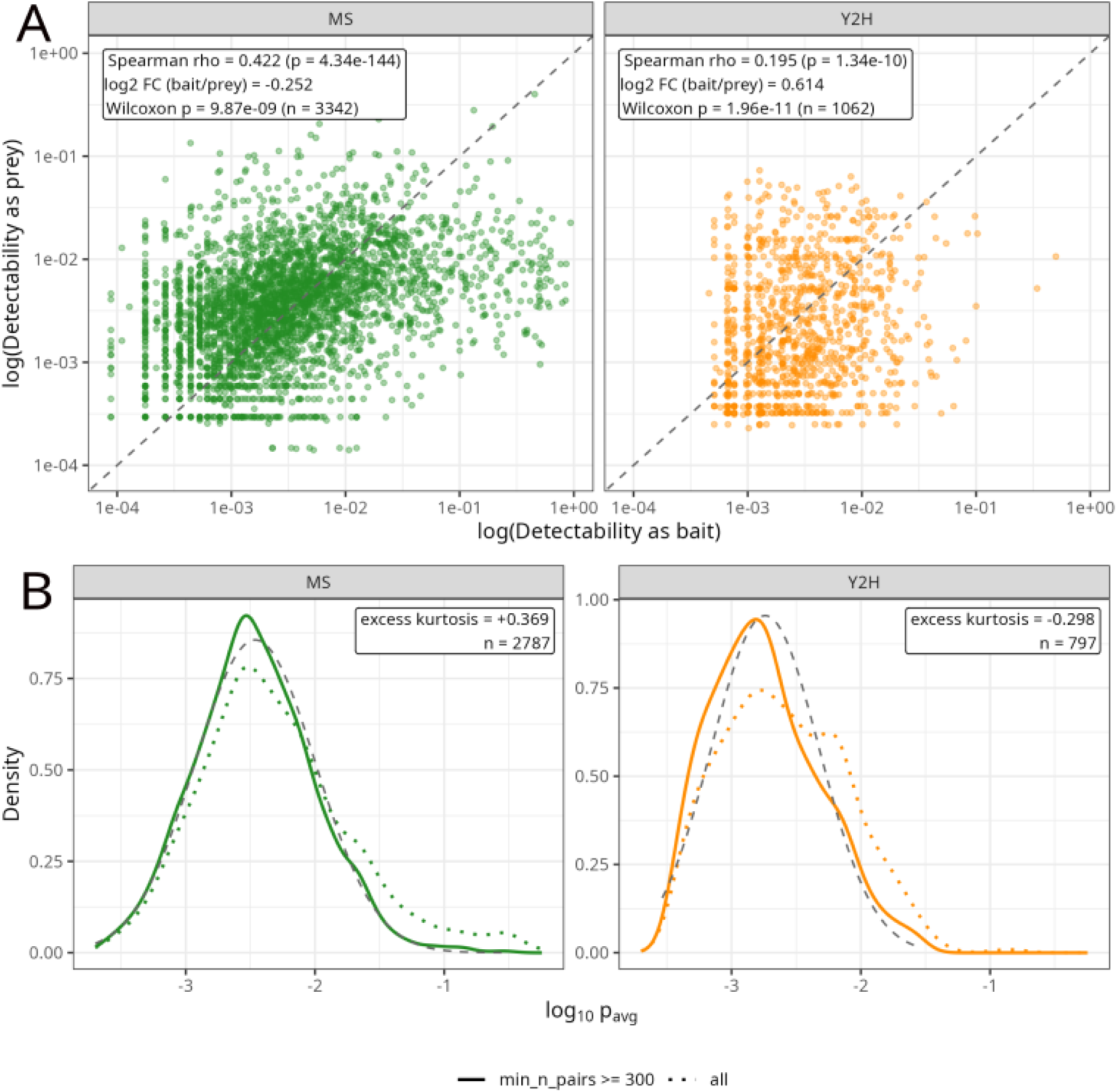
Protein detectability is role-dependent but concordant across roles, and approximately log-normal once sparsely tested proteins are excluded. (A) Per-protein detectability as bait versus as prey, for proteins with at least two tested pairs in each role, in the MS (left, green) and Y2H (right, orange) datasets; dashed line is the identity. The two role-specific estimates are moderately correlated in MS (Spearman ρ = 0.422, n = 3,342) and in Y2H (ρ = 0.195, n = 1,062). Detection is on average higher as prey in MS (log_2_ FC bait/prey = −0.252, Wilcoxon p = 9.9 × 10^−9^) and higher as bait in Y2H (log_2_ FC = 0.614, p = 2.0 × 10^−11^). (B) Density of the role-adjusted average detectability, for all proteins (dotted) and for proteins tested against at least 300 partners (solid), with a fitted normal reference (grey dashed). Restricting to well-tested proteins removes the upper tail present in the unfiltered distribution, leaving excess kurtosis of +0.369 (MS, n = 2,787) and −0.298 (Y2H, n = 797)

We would expect the detectability of a protein tested against a sufficient number of other proteins to be representative of a protein’s number of interactions, or its degree. Therefore we argue that the shape of the *p_avg_* distribution is telling of the human protein degree distribution as a whole. Therefore, we evaluated excess kurtosis of a log-normal fit on protein detectability tested against a minimum of 300 proteins (**Figure 5B**) observing the lack of heavy tail present on the unfiltered data. This means that the heavy tail in our data is best explained by repeat studies containing single or few baits, where no non-interaction data could be inferred.

### Training and evaluating machine learning of PPIs

The task of protein-protein prediction is hard and machine learning approaches to this problem face multiple challenges. The first is lightweight and can be performed with minimal computational effort. While initial studies reported good performance, more recent studies questioned good performance due to data leakage, non-disjoint sets of proteins or unmatched positive-negative degree distributions in train/validation [16]. When properly adjusting for these factors, the prediction accuracy drops close to random. On the other hand, structure based approaches like AlphaFold multimer or RosettaFold report stellar performance [15,40]. None of those approaches are trained or evaluated on experimental negative PPI data yet. Given the high potential of those methods to complete the interactome we suggest incorporating the negative PPIs for training and performance estimation. As retraining of well performing structural methods is beyond our computational scope, we just demonstrate here, how negative data leads to different prediction results on sequence based methods and illustrate how performance estimates incorporating negative interactions change even for high performance methods.

Given the high computational demand of structure based methods and low performance of sequence based ones, we first ask if validated negative data impacts the identity of predicted interactions. We set up a workflow as in (**Figure S1**). In essence, the interaction data were subsetted by different replication thresholds and degree-balanced against thresholded subsets of experimental non-interaction data. Each dataset (Combined, MS, Y2H) were validated and tested on the same subset, while each threshold-configuration training set was balanced to an equal number of edges (max 5% deviation). Finally, each train set had ∼5% of edges randomly removed and rebalanced in order to form 10 subsets with 90-95 % overlap. All training data are degree-balanced and all splits (train/validation and test) were node-distjoint to prevent leakage.

For each of the 10 subsets per threshold-configuration training sets and the three test sets, non-observed (NO) random negative sets were obtained from the complement of the positive set (**Method 4**).This resulted in high data overlap threshold-configuration subset experimental negative data and a very low overlap between their NO negative data counterpart. Due to the downsampling to equal number of edges between experimental threshold-configurations the data overlap was low (**Figure S2**). This setup allows us to estimate the effect of training experimental negative data compared to non-observed interactions.

While as expected [16]performance between models trained on the experimental negative data versus non observed was similarly low (**Figure A/4B**,**Figure S3**), significant differences in the Jaccard index of HRNI set predictions between HRNI trained model and a NO-trained model (median: 0.59-0.74) compared to NO-trained models (median:0.61-0.76) (**Figure 4C**) were observed. This demonstrates that using the here defined set of likely non-interacting protein pairs affects prediction results when used as negative training data.

To allow researchers to further explore the impact of our negative data on training models under fair training conditions [17], we provide balanced splits containing both experimental and non-observed data available at the PPI database HIPPIE [11].

**Figure 4:**
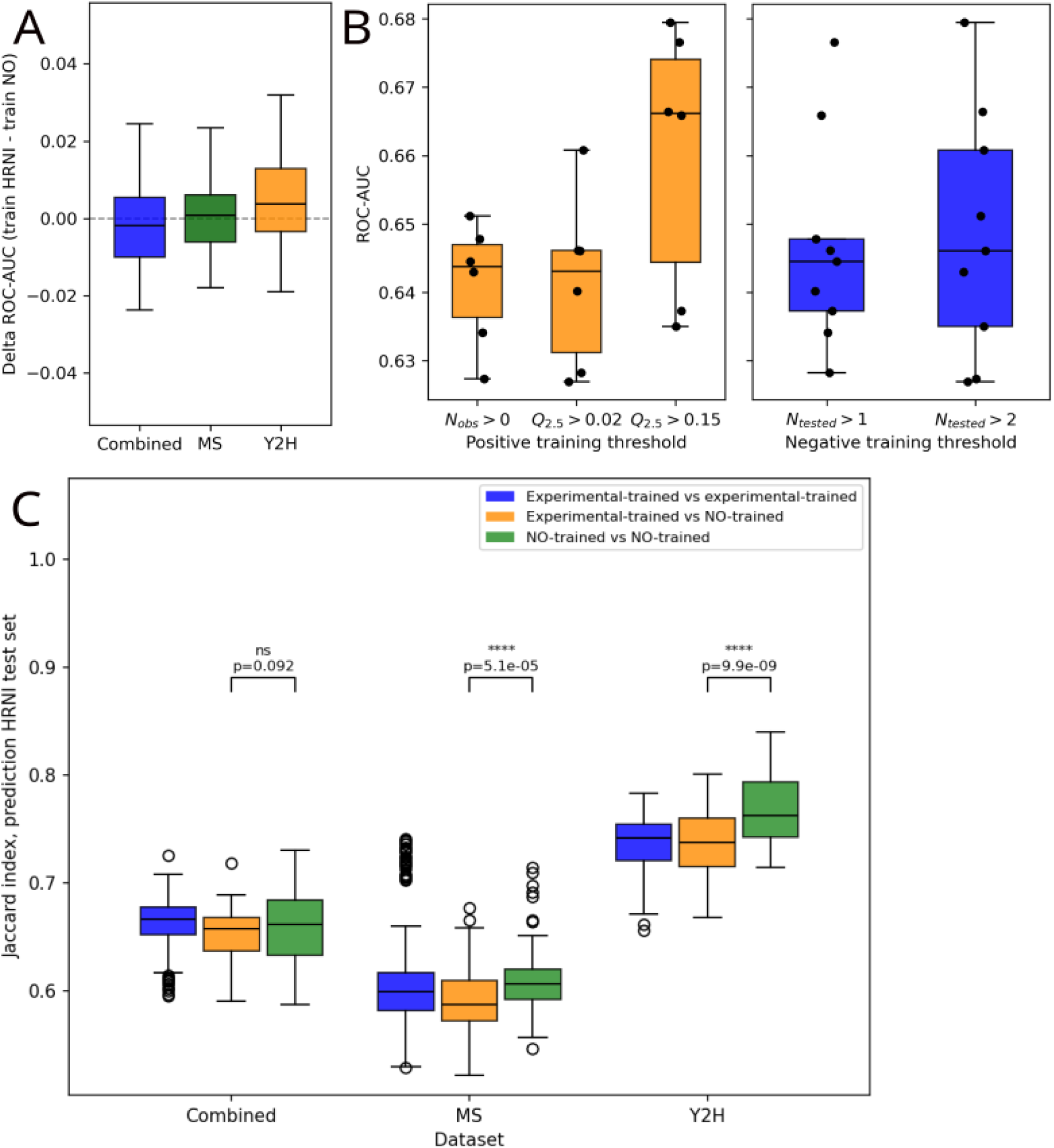
XGBoost classifiers were trained on concatenated mean-pooled ESM-2 embeddings for each protein pair and provided with feature vectors in both directions. (A->B, B->A) for all three detection datasets (Combined, MS and Y2H). Positive data were defined via replication thresholds (*Q*_2.5_ > [0.02, 0.15], or *N_observed_* > 0) paired to a thresholded experimental negative data (*N_tested_* ≥ [1, 2], *N_observed_* = 0). Each threshold configuration was subsampled into 10 partially overlapping positive sets (90–95% shared pairs), each paired with its corresponding negative set and degree-balanced. For each subsets an degree-equivivalent non-observed random set was generated. Within a detection dataset, all models shared the same balanced validation set and were evaluated on the same held-out HRI/HRNI test set. (A) Delta ROCAUC is the difference between the models trained on experimental or non-observed negative data. The boxes show the distribution between the 60 subsamples datasets between per detection dataset. (B) Due to the high data overlap between the random subsampled models, the graph shows mean performance between the subsets. Each dot represents the mean of means across data-configurations and detection dataset, following negative and positive replication thresholds. (C) Negative prediction concordance, between HRNI-HRNI models, HRNI to their NO counterparts and between NO trained models. HRNI models had less overlap in prediction than NO models, even though the HRNI subsampled sets are highly similar while NO sets are highly dissimilar (**Figure S2**).

## Discussion

Protein-protein interaction networks underpin much of our understanding of cellular organisation as they define the repertoire of reactions available to a cell, the modules through which signals propagate and the routes by which alterations translate into disease. Subsequently, accurate representation in the data is imperative for any subsequent analyses as biases, false positives and negatives might muddle biological meaningful conclusions. At the same time, the vast majority of information generated by experiments is not reported: studies and PPI databases report interactions that were detected, but largely discard the experimental search space in which they were identified, i.e. they do not report which proteins have been tested but do not interact. The central claim of this work is that much of this information can be recovered. Starting from approximately 0.5 million reported interactions, we reconstructed evidence for 97 million tested protein pairs corresponding to approximately 182 million individual tests. This provides information on both sides of the interactome: repeated observation frequency strengthens evidence for an interaction, whereas repeated testing without detection provides evidence for a non-interaction. The identification of such a test space, and subsequent aggregation, have many important and interesting applications. We showcase five such applications: (i) FDR/FOR estimation of future interaction identification experiments, (ii) use as a benchmarking tool and calibration of computational PPI prediction (iii) role-specific detectability for AP-MS and Y2H, (iv) insights into the shape of the human-interactome, and (v) use as experimentally supported negative training data in machine learning applications.

We argue that such an experimentally defined search space, and in particular the resulting set of highly likely non-interacting protein pairs, can help address several problems in the network biology field. We rather illustrate the potential of the reconstructed search space here than fully leverage the negative data ourselves, as we believe that many of these applications should ultimately be a community effort. Systematic model development and benchmarking, as well as experimental calibration, are substantial and time– and resource-intensive efforts in their own right. We therefore see this work as both providing a roadmap/blueprint towards a more accurate completion of the human interactome and making the underlying positive and negative experimental evidence accessible as a resource for the community.

Instead, we leverage the now extensive experimental PPI literature to derive negative evidence directly from experimental search space. Completion of the human PPI network will nevertheless require a major effort. Our work can contribute to both computational and experimental charting of its missing pieces. Importantly, this is not only a question of what has not yet been measured: current networks are also distorted by excessive study of particular proteins and by technical biases favouring their inclusion or detection. Our negative data allows us to estimate rather what proteins that each bait have been tested against and how often? With the high FNR reported in literature [6,9] and estimated by us, repeated testing of these interactions are necessary for calling an interaction. With this framework we can also estimate how many proteins a bait has been tested against, which lets us estimate a protein’s relative interaction rate, providing additional information on the proteins role in the interactome. However, this also allows us to evaluate the specificness of a protein’s interactions as either bait or prey, giving us a way to rank all proteins’ stickiness or flag possible auto-activators. In turn this could be employed for cleaner readouts of future studies. Additionally, from the detectability of proteins we might glean the shape of the true degree distribution. The current question is, does its power-law nature come from biology or the aggregation of experimental bias? Our data seems to support something else than the power-law. When only considering proteins with a sufficiently large aggregated search-space, the distribution resembles more that of a log-normal distribution than a power-law. However, this comes with some caveats: namely, the distribution we see might rather be the degree-distribution of proteins detected in MS/Y2H. Secondly, any potential “true” non-interacting proteins are missing from the data as, paradoxically, they will never provide negative data under this approach.

Our results with Xgboost sequence-embedding models show both promise and limitations. A limitation as we could not improve the predictor performance but a promise as these results risk reflecting bias than true signal. Nevertheless, we see and decrease in prediction agreement on negative data compared among HRNI trained models compared to non-observed models for Y2H and MS, even though the the data overlap the HRNI training sets are very high (Jaccard 0.9-0.95) and the NO training sets are practically zero. This implies that the small difference in the HRNI data somehow impacts the heterogeneity in prediction of the model more than non-similar NO data. As of yet we do not know what is driving this heterogeneity, but the datasets containing balanced training sets on HRNI data are available at https://hippie-db.net. For stricter HRI sets we observe a notion of an increase in performance with stricter thresholds but are underpowered to test it, one possible explanation of this is the increased coevolution among the more studied pairs.

However, future charting the unexplored interaction space therefore needs to be accompanied by identifying and reducing these biases. The search-space framework is detection method agnostic, as long as the experiment ensures that each protein detected is part of the same search-space, could be extended to other detection methods such as protein proximity labeling, or protein crosslinking methods. An additional extension is cellular context, such as cell type or tissue specific interactomes where sufficient PPI and cell type data exists. In the end, the recovering of both positive and negative experimental evidence provides a step towards a more complete and less biased human interactome.

## Methods

### 1. Estimation of pair-wise probability of detection

IntAct human PSI-MITAB (released 2026-01-14) and were filtered for human proteins and interactions detected by selected methods (**Table 2**). Canonical isoforms were identified via UniProt API, non canonical isoforms were and proteins without sequences available in Uniprot were discarded. Self-interactions were filtered out since general AP protocols cannot detect them. Single studies were identified via the combination of pubmed id and detection method. Finally, any single study reporting a single interaction were dropped as we hypothesised that these studies were hypothesis-based around that single interaction and not providing quantitative interaction data.

The search-space of each single study was identified via the prey identified. The baits were not considered part of the search space as these are of the artificially transducted and therefore not guaranteed to be available to all other baits. From this search space the non-interaction data was inferred for each experiment. The bait-prey observed/tested counts were subsequently aggregated from all available experiments and subsequently aggregated on a protein pair basis, removing the bait-prey directionality of the data resulting in 97 M unique protein pairs constituting 182 M tests.

To estimate the probability of detection we fit the observed values to a beta-binomial distribution as:

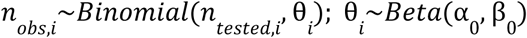

Where *i* denotes the specific protein-pair. The empirical prior was defined on the global detection ratio *p* = *N_obs_* /*N_tested_*, with α_0_ = *kp* and β = *k*(1 − *p*), where *k* = 1 is the pseudo-count meaning that the prior strengths were set at the strength equal to a single experiment. This prior was set as the interaction rate within the full human search-space is very low, and this prior reflects that fact.

For each interaction the posterior probability of detection were calculated as:

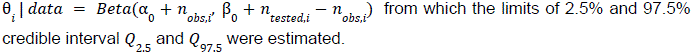

Due to the very low prior sets of HRNI are reported by minimum number of times the protein pair has been tested and never reported an interaction rather than by a threshold on *Q*_97.5_.

### 2. Co-annotation analysis HRNI vs HRI

Go-BP annotations were obtained from the full genome using the MyGene [45] python module (obtained 20-08-26) and sub-cellular localisation data from Human Protein Atlas [30] (obtained 6-04-26). The subcellular annotations were filtered on reliability: *Supported, Approved* or *Enhanced* and only the *Main location* annotation were used. Additionally, all annotations were filtered to a minimum of 400 genes per term. This was done due to the majority of GO-BP terms being annotated to few genes with a high protein interaction connectivity. The small number of genes limits the number of negative interactions that could be tested within the set of proteins annotated. By selecting the larger annotation specific gene sets we can both lessen the impact of multiple hypotheses correction and decrease the influence of smaller annotation sets. We the OR as 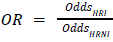 computed the proportion *p* of edges sharing that annotation for the HRI and HRNI datasets and the odds for each dataset *Odds* = *p*/(1 − *p*). The annotations are properties by the proteins (nodes) and not the protein pairs (edges) edges sharing a protein cannot be considered independent. Therefore, we assessed the uncertainty of OR estimates using node-level bootstrapping. However, shared annotation is a property of the edges so to get a protein level statistic we calculate the rate of shared annotation per protein: the total number of shared annotation edges with protein *i* is *N* and the total number of edges containing protein *i* is *N_i_*. This gives us the protein annotation counts and non-annotation counts which we uniformly sample with resampling. For each sampling the the HRI and HRNI rate can be estimated as *p* = Σ*N_s_*_,*i*_/Σ*N_i_*. We conducted 5000 such resamplings and estimated the 95% CI from the interval. Additionally we report two-tailed p-values were estimated using said samplings and q-values adjusted using Benjamini-Hochberg (**Table S3**).

### 4. ML-splits strategy and model setup

The validation set was degree-balanced between HRIs and HRNIs using an alternating max-flow approach (**Method S1**), allowing for a maximum of 5% total degree deviation while minimizing the percent individual protein deviation (**Figure S1**). Nodes exclusive to either the HRI or HRNI set were dropped from the validation set and reassigned to the test set. To give the validation and test sets roughly equal numbers of positive edges, we then selected a random subset of nodes and refined the split by reassigning nodes between the two sets over 20 iterations, selecting the iteration with the highest sample balance.

To quantify the contribution of validated non-interactions across thresholds, we built degree-balanced training sets, each disjoint from the test and validation sets. For every positive/negative threshold configuration (**Figure 4B**), training edges were drawn from the data remaining after the test/validation split and balanced to a common size (**Method S2**). Each training set was further downsampled by 5 %, repeated ten times with different random seeds, to generate replicate sets for estimating the stability of PPI-prediction performance in downstream evaluation.

In order to obtain a representative non-observed negative data for each downsized degree balanced set. The complement of the positive protein pairs were balanced against the positive edges.

### Classification setup and protein-embeddings

For sequence information, the canonical transcripts were retrieved from Uniprot. Protein sequence embeddings were generated using the ESM-2 model (esm2_t33_650M_UR50D), deployed via Hugging Face’s Docker image (transformers-all-latest-torch-nightly-gpu [04–2026]).

For each protein, the final representation was computed by taking the mean across all embedding dimensions, resulting in a 1,280-dimensional feature vector. For each protein pair, the interactions were represented both as protein A –> protein B and protein B –> protein B. The features of each interaction were concatenated, producing a 2,560-dimensional x 2 input matrix per edge.

The model selected was an XGBClassifier implemented in XGBoost(ref). Hyperparameters were optimized over 5 iterations using Bayesian optimization (parameter range in **Table 2**) with expected improvement, minimizing cross-entropy against the validation set. The final model was trained on both train and validation, over 100 estimators, and a gamma-value of zero with the default tree-based boosting (gbtree).

**Table 2:** Hyperparameter space considered for XGBClassifier.

| Parameter | Range |
| --- | --- |
| max_depth | 3 - 10 |
| learning_rate | 0.01 - 0.3 |
| min_child_weight | 1 - 50 |
| colsample_bytree | 0.3 - 1.0 |
| subsample | 0.5 - 1.0 |

### Implementation

The full workflow is implemented as a Snakemake workflow for data formatting, search-space inference, downstream analyses and PPI classification benchmarking. Data handling is mostly written in python [3.12]. Version control is achieved through git and environment control through Conda and apptainer. GLMM fitting was implemented using the MixedModel [v.5.9.0] julia module. Plotting is mostly done in R.

The computationally heavy step of embedding computations were done on the IEO HPC using a Nvidia H200 GPU using Hugging Face’s *transformers-all-latest-gpu* container.

Remaining analysis was conducted on a 48-core Intel(R) Xeon(R) Silver 4214 CPU 188 GB of ram.

Full workflow and list of software versions are found at https://github.com/JoelAAs/PPI-bias (main,7bbe392)

## Supplement

### Supplemental methods

**Method S1:** *Degree balancing by alternating max-flow*

To stop node degree from acting as a predictive shortcut, we subsample the interaction network *G*_+_ and non-interaction network *G*_−_ so they share the same per-protein degree and edge count. Both networks are first restricted to their shared proteins, repeating the pruning until the node sets match so every protein has at least one positive and one negative edge.

To match *G*_+_ to the degree of *G*_−_, we select a degree-bounded subgraph via max-flow. Each protein *v* is split into a source-side and a sink-side copy, both connected to the source/sink with capacity equal to its target degree *d*(*v*), initial network set to degree in *G*_+_. Every candidate edge (*u*, *v*) is added as two flow-edges, (*u*, *v*) and (*v*, *u*). Push-relabel max-flow (graph-tool) saturates as many as the capacities allow, selecting the largest set of edges whose degree sequence does not exceed twice target degree (once for source, once for sink). To reconstruct the selected graph: All pair with both flow-edges saturated, (*u*, *v*) and (*v*, *u*), are kept. Any pairs with only one edge saturated are resolved greedily by processing the largest remaining slack first and adding (*u*, *v*) only while both proteins have a degree below target. The slack is defined as the missing number of degrees per protein after selecting all full-edges. If degree is still not fulfilled and there are remaining but damaging half-edges, edges are added if their as long as they improve the percentual balance, i.e. if the addition of an edge increases the overshot of a degree target of 10% but decreases by 50% for the other node, the edge is added.

Since each pass only removes edges from the graph building the flow-edges, we alternate the target and edge network until both per protein degree converge.

#### Pseudocode

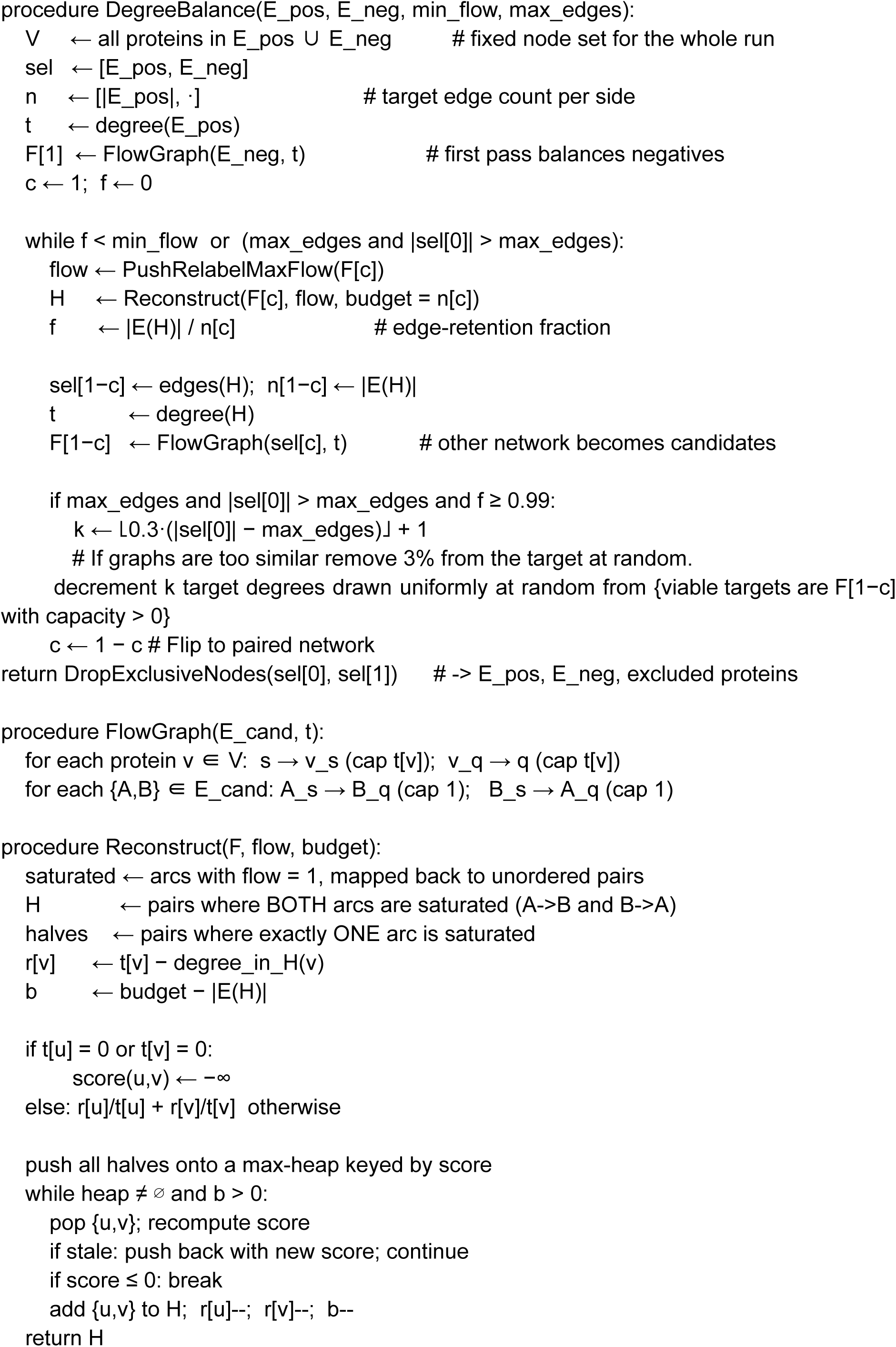

**Method S2:** Pruning and balancing of train sets.

To achieve different training sets with a similar number of edges all train sets (from the same datasets eg. Y2H, MS) was degree-matched between positives and negatives with **Method S1**, then all configurations were equalized to a common edge count within 5%.

#### Pseudocode

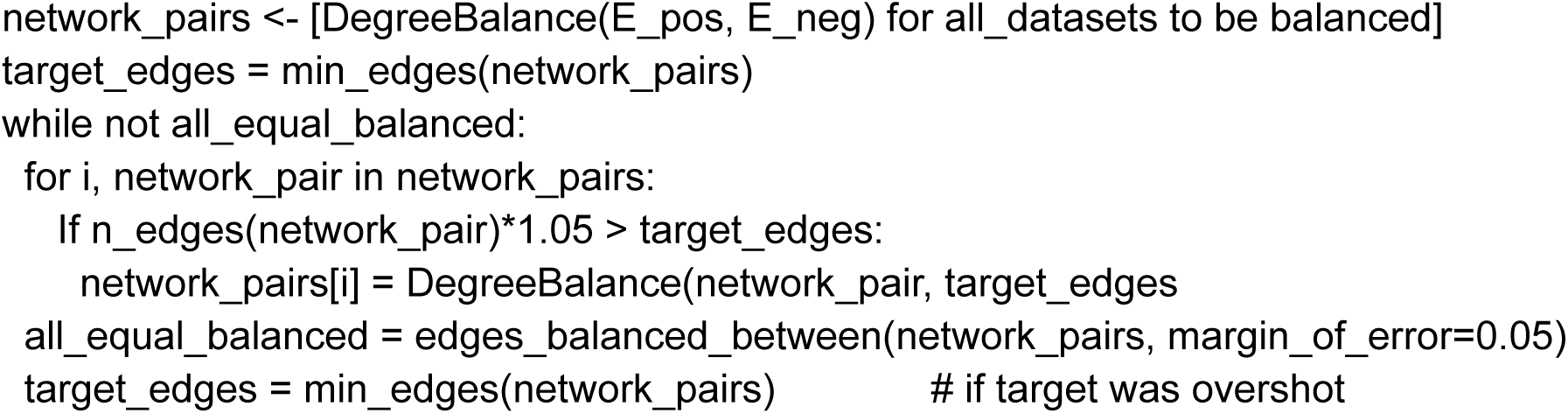

## Supplemental Tables

**Table S1:**
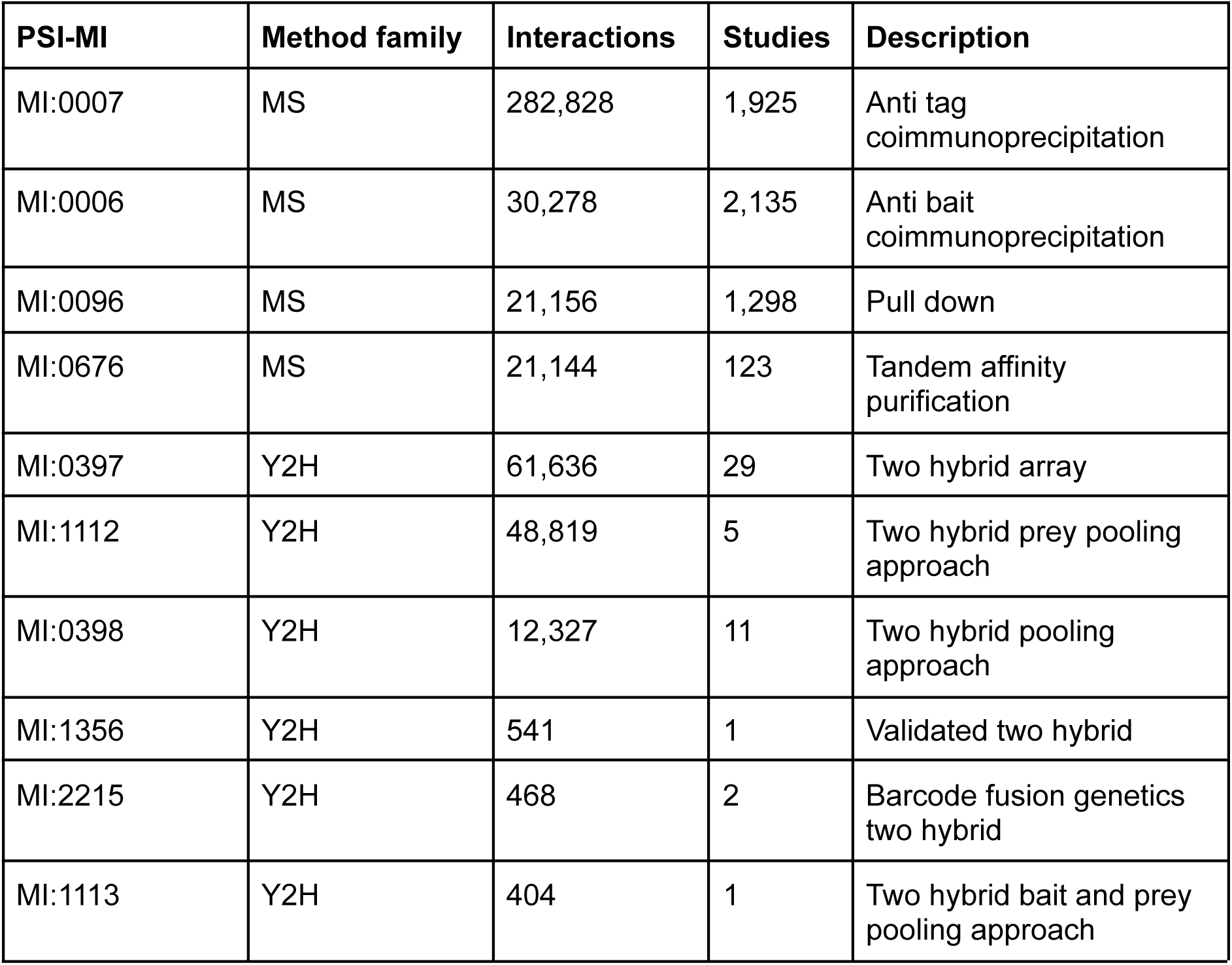
Selected bait-prey methods, number of filtered interactions and studies.

**Table S2:**
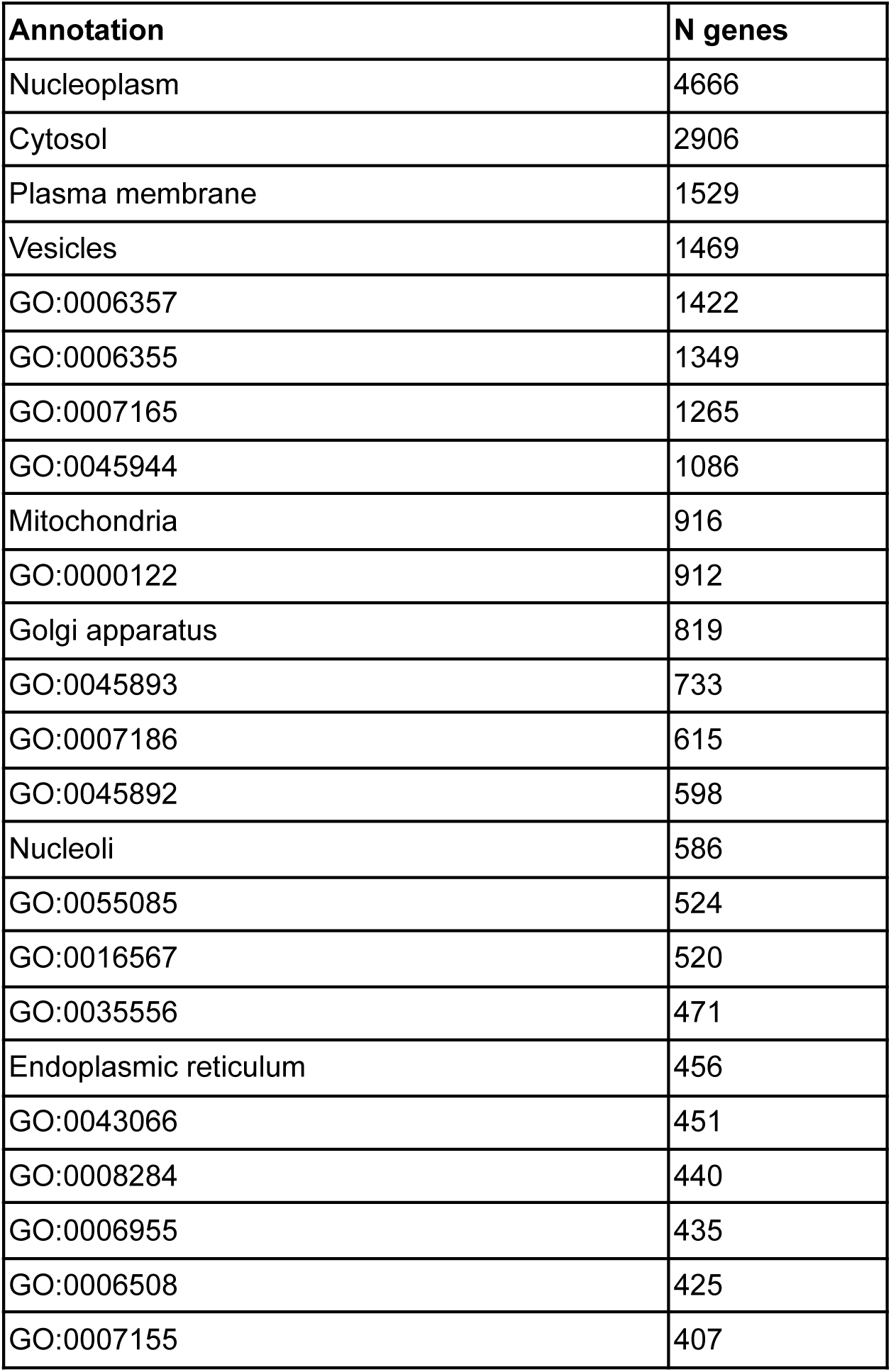
Number of genes associated with GO-process or Localisation.

| Annotation | N genes |
| --- | --- |
| Nucleoplasm | 4666 |
| Cytosol | 2906 |
| Plasma membrane | 1529 |
| Vesicles | 1469 |
| GO:0006357 | 1422 |
| GO:0006355 | 1349 |
| GO:0007165 | 1265 |
| GO:0045944 | 1086 |
| Mitochondria | 916 |
| GO:0000122 | 912 |
| Golgi apparatus | 819 |
| GO:0045893 | 733 |
| GO:0007186 | 615 |
| GO:0045892 | 598 |
| Nucleoli | 586 |
| GO:0055085 | 524 |
| GO:0016567 | 520 |
| GO:0035556 | 471 |
| Endoplasmic reticulum | 456 |
| GO:0043066 | 451 |
| GO:0008284 | 440 |
| GO:0006955 | 435 |
| GO:0006508 | 425 |
| GO:0007155 | 407 |

Table S3:

**Table S3:**
| annotation | OR<br>Combin<br>ed | q<br>Combin<br>ed | OR<br>MS | q<br>MS | OR<br>Y2H | q<br>Y2H |
| --- | --- | --- | --- | --- | --- | --- |
| GO:0000122 (negative regulation of transcription by RNA polymerase II) | 12.776 | 2.40E-04 | 18.721 | 2.29E-04 | 3.212 | 4.80E-04 |
| GO:0006355 (regulation of DNA-templated transcription) | 6.257 | 2.40E-04 | 11.187 | 2.29E-04 | 1.836 | 4.80E-04 |
| GO:0006357 (regulation of transcription by RNA polymerase II) | 9.171 | 2.40E-04 | 18.46 | 2.29E-04 | 1.633 | 8.73E-04 |
| GO:0006508 (proteolysis) | 0.46 | 1.39E-01 | 0.426 | 9.88E-02 | 0 | 4.80E-04 |
| GO:0006955 (immune response) | 6.278 | 2.40E-04 | 8.476 | 2.29E-04 | 0 | 4.80E-04 |
| GO:0007155 (cell adhesion) | 9.042 | 2.40E-04 | 10.191 | 2.29E-04 | 2.442 | 3.67E-01 |
| GO:0007165 (signal transduction) | 5.181 | 2.40E-04 | 6.641 | 2.29E-04 | 2.186 | 4.80E-04 |
| GO:0007186 (G protein-coupled receptor signaling pathway) | 6.365 | 2.40E-04 | 11.238 | 2.29E-04 | 0 | 4.80E-04 |
| GO:0008284 (positive regulation of cell population proliferation) | 6.914 | 2.40E-04 | 8.383 | 2.29E-04 | 2.265 | 3.21E-01 |
| GO:0016567 (protein ubiquitination) | 17.932 | 2.40E-04 | 22.386 | 2.29E-04 | 7.59 | 4.80E-04 |
| GO:0035556 (intracellular signal transduction) | 5.684 | 2.40E-04 | 5.927 | 2.29E-04 | 2.992 | 6.30E-02 |
| GO:0043066 (negative regulation of apoptotic process) | 6.874 | 2.40E-04 | 9.02 | 2.29E-04 | 1.597 | 3.38E-01 |
| GO:0045892 (negative regulation of DNA-templated transcription) | 17.269 | 2.40E-04 | 24.474 | 2.29E-04 | 3.097 | 4.80E-04 |
| GO:0045893 (positive regulation of DNA-templated transcription) | 16.104 | 2.40E-04 | 22.501 | 2.29E-04 | 3.022 | 4.80E-04 |
| GO:0045944 (positive regulation of transcription by RNA polymerase II) | 8.683 | 2.40E-04 | 11.58 | 2.29E-04 | 1.653 | 1.85E-02 |
| GO:0055085 (transmembrane transport) | 3.442 | 2.62E-03 | 5.051 | 2.09E-03 | 1.773 | 7.32E-01 |
| Cytosol | 1.937 | 2.40E-04 | 2.256 | 2.29E-04 | 1.079 | 3.84E-01 |
| Endoplasmic reticulum | 3.083 | 2.40E-04 | 3.069 | 2.29E-04 | 0.886 | 7.86E-01 |
| Golgi apparatus | 1.952 | 2.62E-03 | 2.061 | 2.09E-03 | 1.445 | 3.67E-01 |
| Mitochondria | 3.738 | 2.40E-04 | 4.475 | 2.29E-04 | 0.524 | 6.30E-02 |
| Nucleoli | 3.7 | 2.40E-04 | 4.49 | 2.29E-04 | 0.255 | 4.80E-03 |
| Nucleoplasm | 3.021 | 2.40E-04 | 3.967 | 2.29E-04 | 1.449 | 4.80E-04 |
| Plasma membrane | 1.899 | 2.40E-04 | 2.312 | 2.29E-04 | 0.537 | 5.17E-03 |
| Vesicles | 1.392 | 1.50E-02 | 1.752 | 2.29E-04 | 0.987 | 9.21E-01 |

## Supplemental Tables

**Figure S1:**
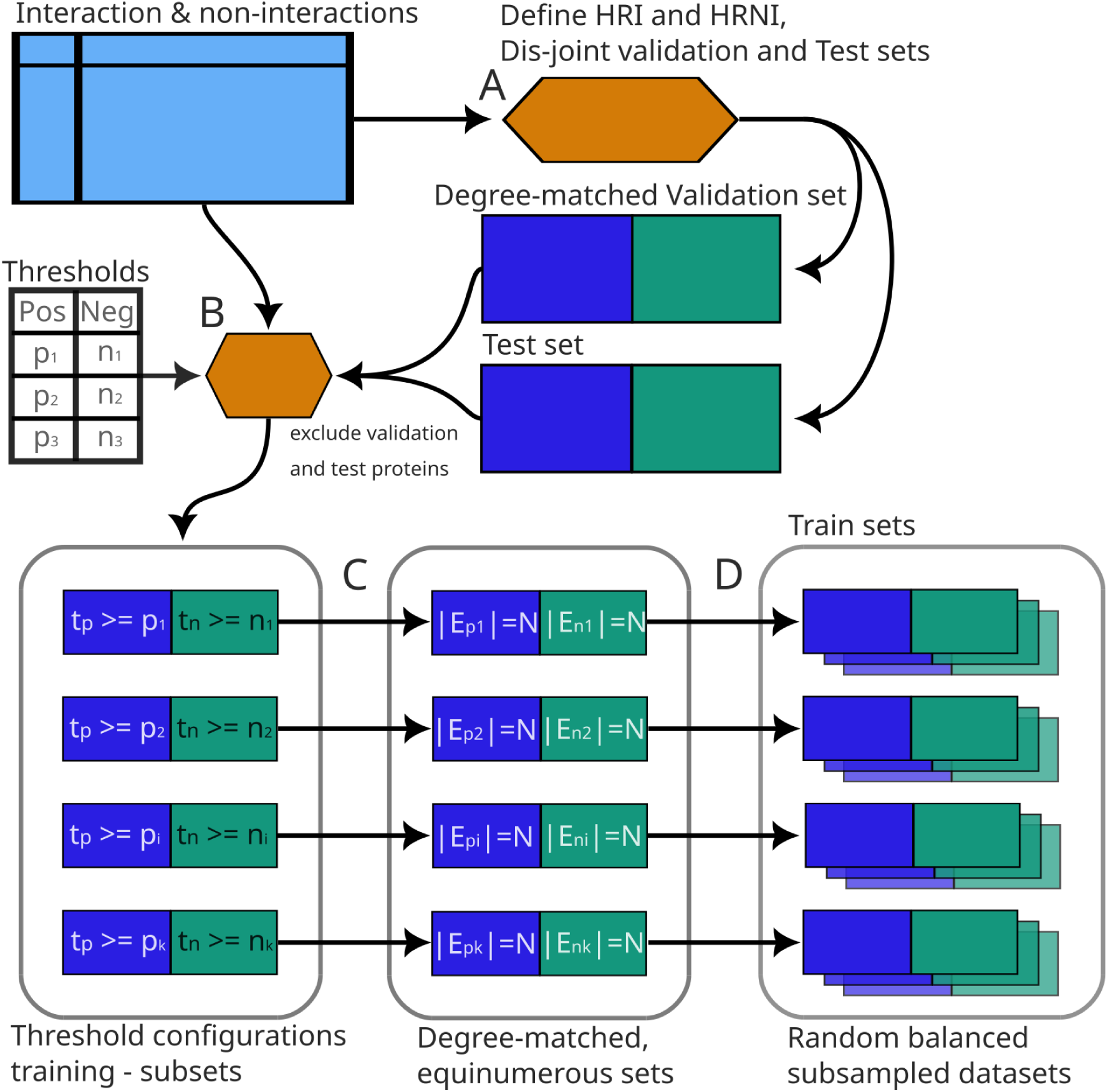
Strategy for creating disjoint, degree-matched train, validation, and test sets. The pooled set of high-replication interactions (HRIs) and high-replication non-interactions (HRNIs): (A) The data are partitioned into a test set and a validation set that share no proteins and hold roughly equal numbers of HRIs. The validation set is degree-matched for positive (interaction) and negative (non-interaction) edges. (B) Unbalanced training sets are then defined for every combination of a positive replication threshold (pᵢ) and a negative replication threshold (nᵢ). Any protein appearing in the validation or test set is excluded, keeping the training data disjoint from the evaluation data. (C) Each threshold-configuration training set is degree-balanced and subsampled to a common edge count N, so all configurations contain the same number of positive and negative edges, allowing for a 5% deviation. (D) Each threshold-configuration set is resampled 10 times with different random seeds; in each replicate, edges are iteratively dropped and the set re-balanced until 5% of the original edges have been removed. This yields 10 random, balanced, degree-matched, equinumerous training sets per threshold configuration.

**Figure S2:**
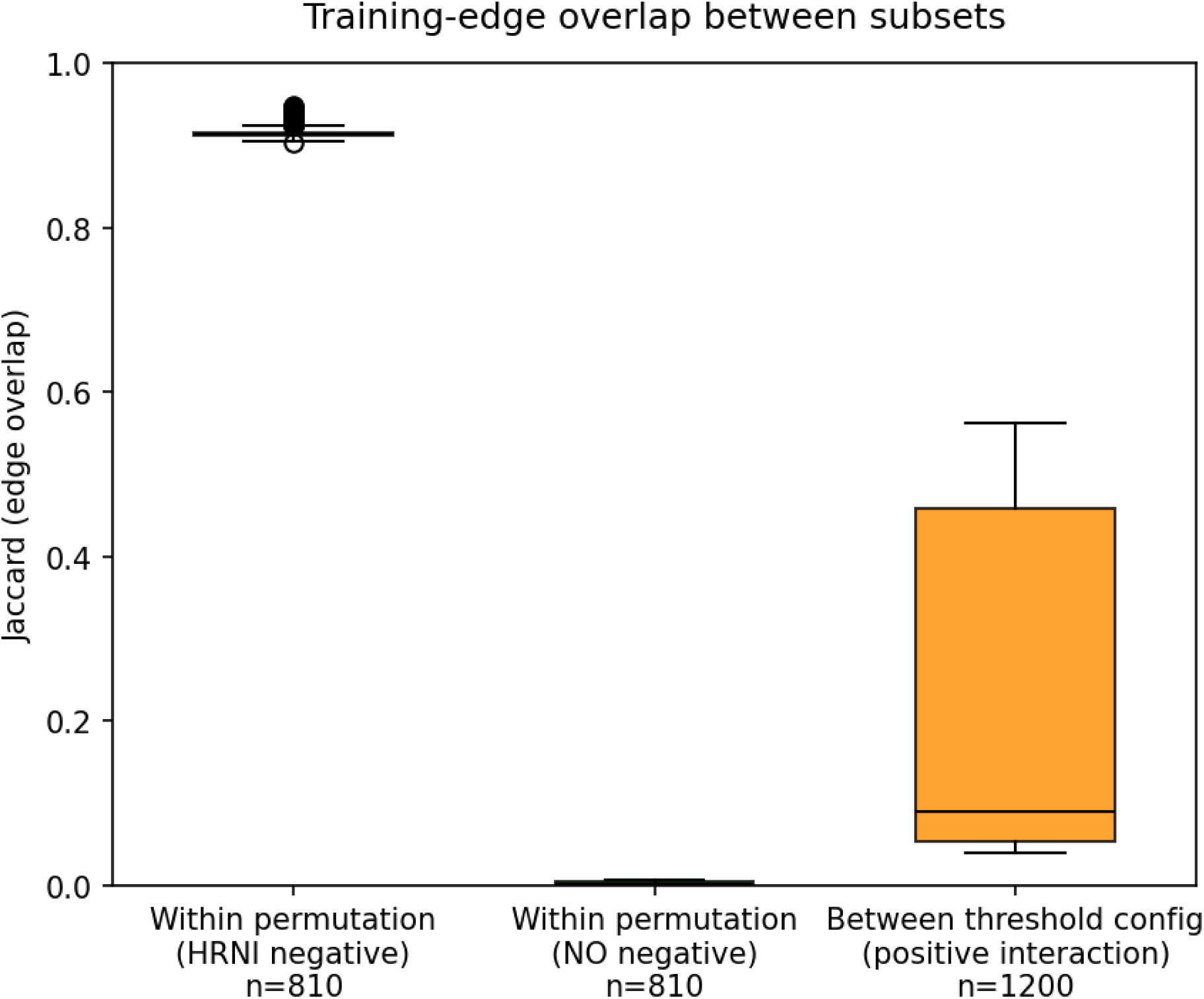
Jaccard index, between and within configuration. The high Jaccard index between negative data in the subsampled training sets, the low Jaccard index between their non-observed observed counterparts and the medium to low Jaccard index between interaction data configuration thresholds form the same detection dataset (due to subsampling to equinumerous edges).

**Figure S3:**
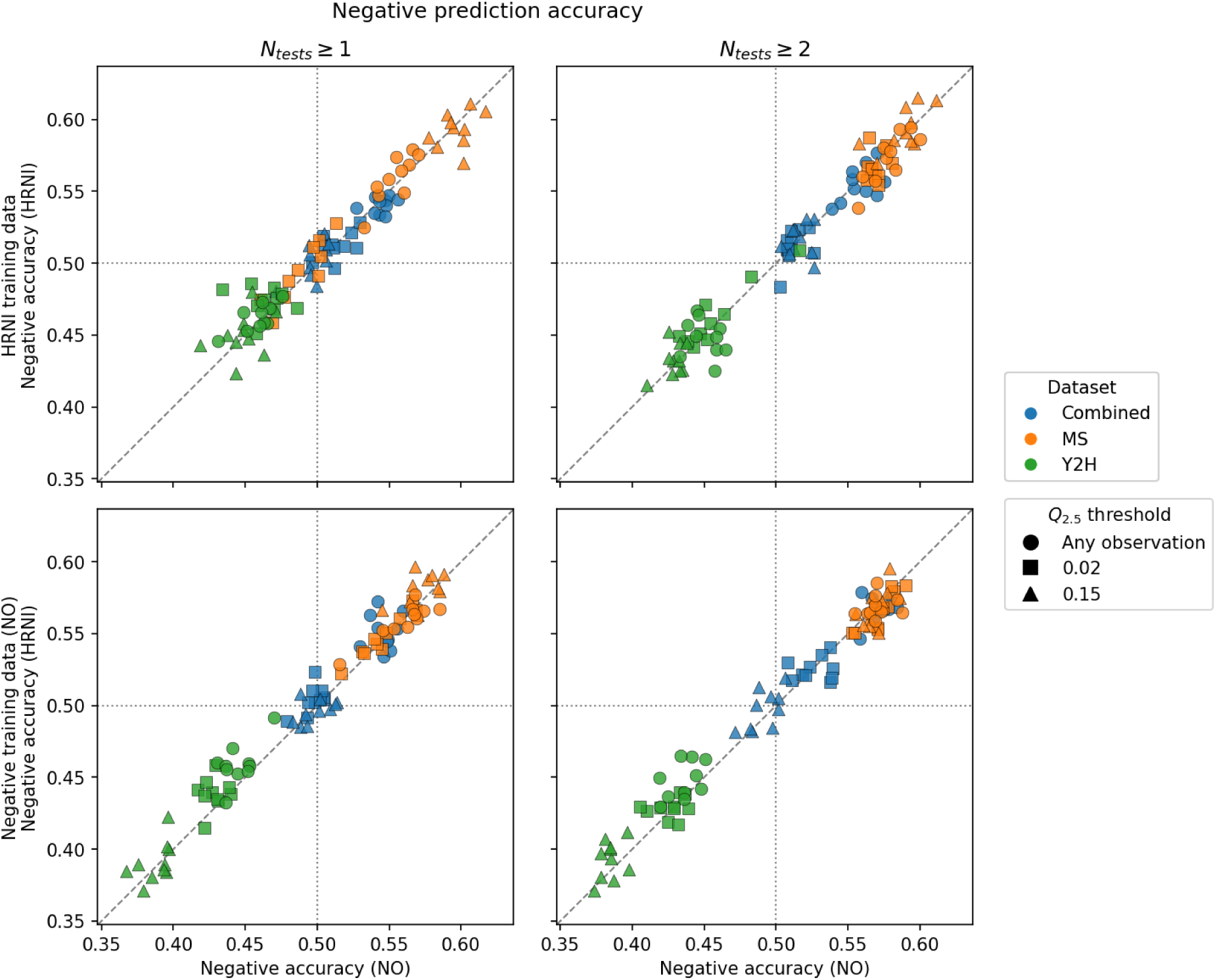
Equal negative accuracy of non-observed or HRNI test data. As evident the models perform equally when tested against experimental HRNI data as equivalent non-observed data, regardless if trained HRNI or non-observed data. This means that the models learn primarily through the constituency of the proteins in the training set, but not their negative edges.

**Figure S4:**
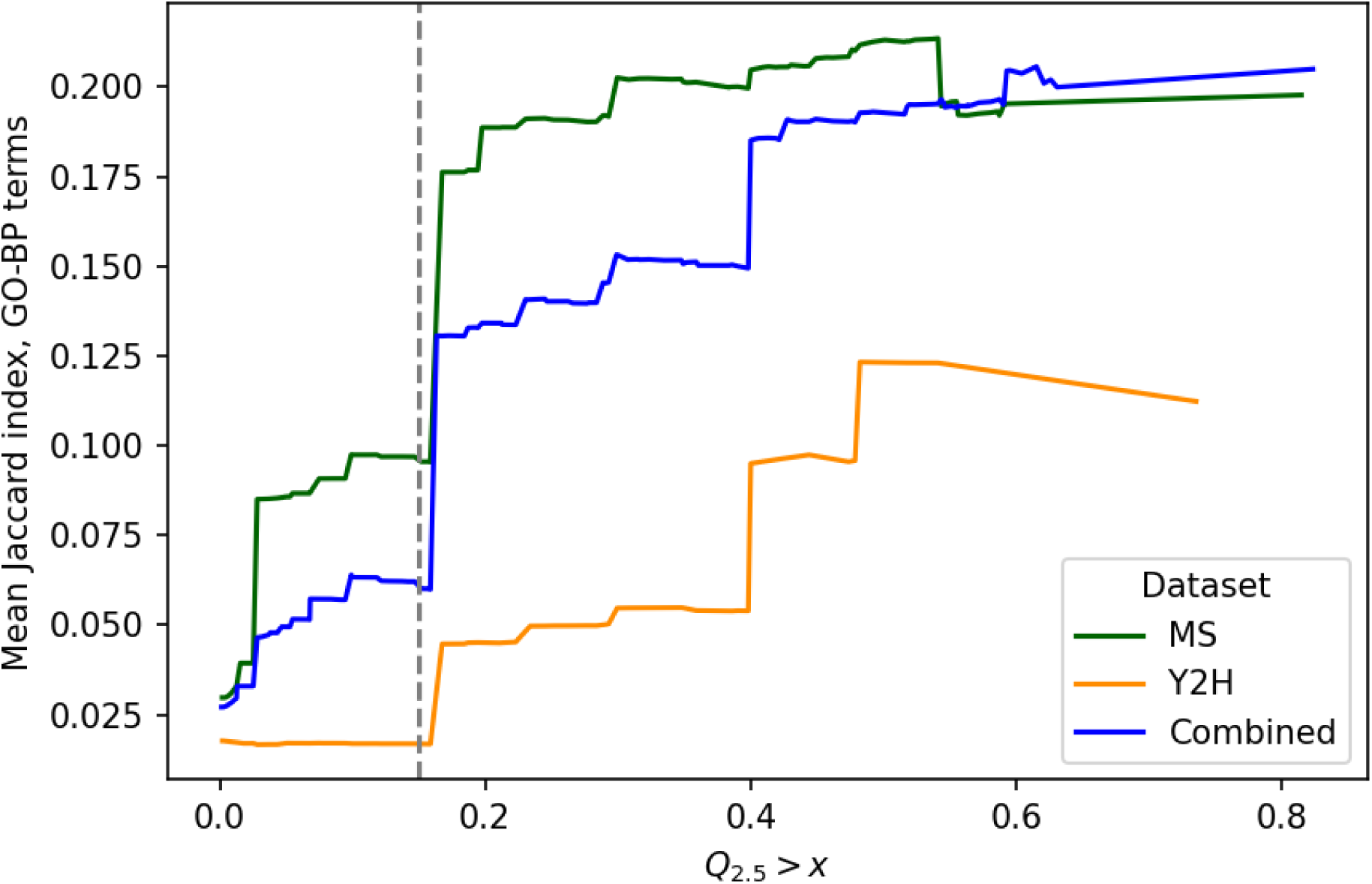
GO-BP Jaccard increases with detectability. The sliding mean jaccard index over lower bound detectability Q.25 > x. As evident by the plot, the higher the detectability estimate of a protein pair is, the higher the propensity of sharing GO biological process terms.

